# Decline in large-seeded species in Danish grasslands over an eight-year period

**DOI:** 10.1101/2021.03.02.433513

**Authors:** Christian Damgaard

## Abstract

A time-series analysis of cover data from 236 Danish grassland sites demonstrated community selection on seed mass. Across four grassland habitat types and during an eight-year period there was a significant decline in large-seeded species. In the analysis, the continuous seed size variable was used to group plant species into functional types. This method was chosen in order to account for the sampling process of the abundance data, and is in contrast to most other analyses of trait selection, where the community weighted mean of the traits is used as the dependent variable. Generally, little is known on the causes for seed size community selection in semi-natural grasslands that are not subjected to successional processes.

## Introduction

In a world where the environment is changing due to climate change and land-use changes, it may be expected that the changing environment may lead to plant trait selection at the community level (*sensu* Vellend 2010), i.e. plant species with favored traits will generally outcompete plant species with other traits. In the past seventy years, there has been a dramatic change in the land-use of grassland habitats, where the use and management of semi-natural grasslands have changed towards intensified agricultural production and fertilization in some areas and abandonment of traditional practices (e.g. cessation of extensive livestock grazing) in others (Poschlod et al. 2005). Concurrently, there has been an increase in atmospheric nitrogen deposition, although nitrogen deposition recently has levelled off and has even shown a decreasing tendency in Denmark (Ellermann et al. 2019). The effects of these land-use changes vary, but, generally, there has been a change in the vegetation towards taller and more competitive plant species (Timmermann et al. 2015), and these land-use changes may also have had an effect on seed mass selection.

Seed size is known to be of general importance in plant life history (e.g. Westoby et al. 2002), and it is of interest to know whether the average seed size is changing across natural habitats in these years as a possible response to the changing environment. In most plant communities, there is large interspecific variation in seed mass, and this variation has been suggested to play a role in a possible trade-off between fecundity and stress tolerance early in plant development (Muller-Landau 2010; Westoby et al. 2002). That is, if a certain amount of resources are allocated to seed production, then the plant species may either produce few heavy seeds or many light seeds and, generally, there is a negative correlation between seed mass and seed number (Coomes and Grubb 2003). Furthermore, there is substantial evidence that seedlings from heavy seeds are more likely to tolerate stressful conditions early in plant development (Coomes and Grubb 2003; Larios et al. 2014; Westoby et al. 2002). Maron et al. (2021) found that the fitness of large seeds increased with competition intensity and that the fitness of small seeds increased with the level of seed predation in a study of 17 forb species in environments with variable competition intensity and rodent seed predation. Generally, seed dispersal distances are relatively independent of seed size (Coomes and Grubb 2003; Thomson et al. 2011), although a positive correlation between seed mass and dispersal distance has been documented when seeds are collected by animals with a hoarding strategy (Jansen et al. 2002). Consequently, the trade-off between seed number (increased fecundity) and seed mass is intricate and depends on the environment, and it is expected that this trade-off may be undergoing selection at the community level (*sensu* Vellend 2010) as a response to changes in the abiotic and biotic environments. However, little work has been done to demonstrate such selection forces in natural habitats that are not subjected to large successional changes.

Recently, there has been increasing awareness that in order to avoid bias in ecological models, it is important to model the sampling variance correctly. This has been demonstrated in model-based ordination (Damgaard et al. 2020; Hui et al. 2015), and more specifically, in the statistical analysis of plant trait changes (Clark 2016; Damgaard 2021). Usually in community ecology, trait selection is investigated using community weighted mean trait values (CWMs) as the dependent variable; where the CWMs are calculated from a fixed species-trait matrix and the measured species abundances in a plot (e.g. Garnier et al. 2016). However, as pointed out by Clark (2016) it is not the CWMs that are sampled, but rather species abundances, and the stochastic properties of CWMs do not arise from variation in traits, but depend on the used sampling protocol to estimate species abundance. One of the ways to solve the problem is to work with discrete plant functional types with known statistical sampling properties as the dependent variable rather than CWMs (Clark 2016; Damgaard 2021).

In this study, I will investigate whether the species composition in four Danish grassland habitats has changed during an eight-year period, in such a way, that the typical seed size has changed, i.e. whether there has been community selection on seed size. This is done by grouping grassland plant species into plant functional types (PFTs) according to their seed mass and investigate whether the proportion among the PFTs has changed across four grassland habitats during the period from 2007 to 2014. This work is a companion study to Damgaard (2021), where the selection forces on the leaf economy spectrum in the same grassland habitats were investigated.

## Materials and Methods

### Sampling design and plant cover data

Hierarchical time-series plant cover data in the eight-year period from 2007 to 2014 from 236 Danish grassland sites were used in the analysis. The data are a subset of the ecological monitoring data collected in the Danish habitat surveillance program NOVANA (Nielsen et al. 2012; Nygaard et al. 2016). All sites included several grassland plots, with a total of 2946 plots that all were resampled three or more times with GPS-uncertainty (< 10 meters) over the sampling years. All plots at a site were either sampled or not sampled in a given year, but not all sites were sampled in a given year. This irregular sampling design results in a considerable among year variation. Including resampling over the years, a total of 8859 vegetation plots were used in the analyses.

All plots were classified as belonging to one of four grassland habitat types: calcareous grasslands (xeric sand calcareous grasslands, EU habitat type: 6120, 99 plots), dry grasslands (semi-natural dry grasslands and scrubland facies on calcareous substrates, EU habitat type: 6210, 1155 plots), acid grasslands (species-rich Nardus grasslands, EU habitat type: 6230, 1129 plots) and wet grasslands (Molinia meadows on calcareous, peaty or clayey-silt laden soils, EU habitat type: 6410, 563 plots). The classification of the habitat types was performed according to the habitat classification system used for the European Habitat Directive (EU 2013; Nygaard et al. 2009). Since the habitat classification was performed for each plot, some sites may contain plots that were classified to different grassland habitat types. Only plots that over the years were consistently classified as belonging to the same grassland habitat type were used in the analyses.

The plant cover, i.e. the relative projected area covered by a species, of all higher plants (Angiosperms) was measured by the pin-point method (Damgaard 2009; Levy and Madden 1933; Lindquist 1931) using a square frame (50 cm X 50 cm) of 16 grid points that were equally spaced by 10 cm (Nielsen et al. 2012). Since the plots were resampled with GPS-uncertainty (< 10 meters), the probability that the location of resampled plots overlapped may be ignored.

In the present study, plant species were aggregated into PFTs according to their seed mass, and one of the advantages of the pin-point method is that it is possible to aggregate taxa at the pin level. For example, if we want to aggregate the cover of two species, then the number of pins that hit either one or both of the species is used as a measure of the aggregated cover of the two species.

At the same time, when the vegetation was monitored it was recorded at each plot whether there were any signs of grazing by animal husbandry, which was used as a possible explaining variable.

### Seed mass

The seed mass values were found in the LEDA trait database (Kleyer et al. 2008), a trait database established from plant measurements in North-West Europe, which is geographically close to the studied sites. There will always be missing values when applying existing trait data sets to national survey data, and while more comprehensive plant trait databases exist, the priority in this study was to use a homogenous and geographically representative trait database in order to avoid data bias.

In order to group the higher plant species into PFTs, the trait values of all the 757 species, where seed mass data were available in the database, were used to group the species into PFTs with similar mean cover: 1: light seeds (log(*seed mass in mg*) < −1), 2: intermediate seeds (−1 ≤ log(*seed mass in mg*)< 0), 3: heavy seeds (log(*seed mass in mg*) > 0), 4: the rest, i.e. species without seed mass data (Fig. 1). The mean site cover of the four different species groups across habitat types and years is shown in Fig. 2.

**Fig. 1.**
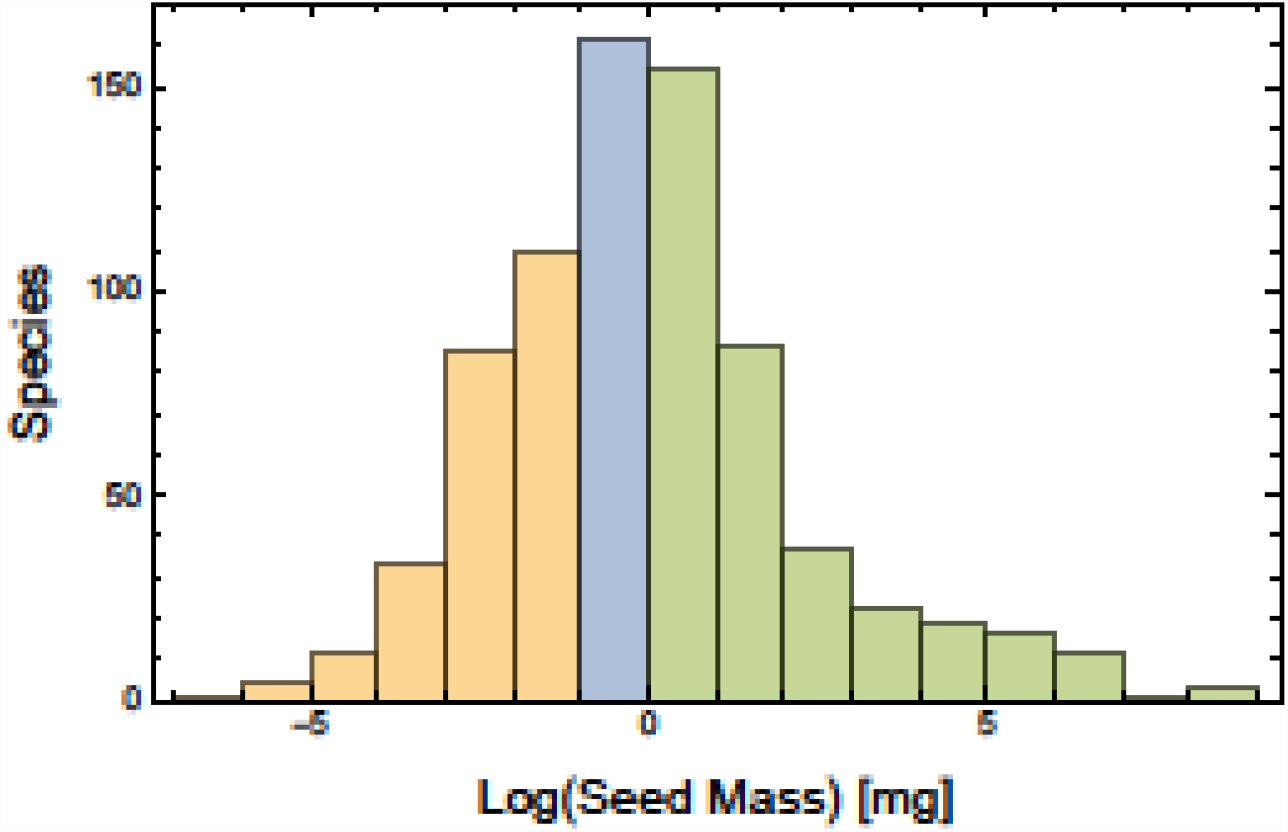
Histogram of seed mass of all the 757 species that was available in the database. The seed mass data were used to group the species into PFTs with similar seed size; yellow: light seeds, blue: intermediate seeds, green: heavy seeds.

**Fig. 2.**
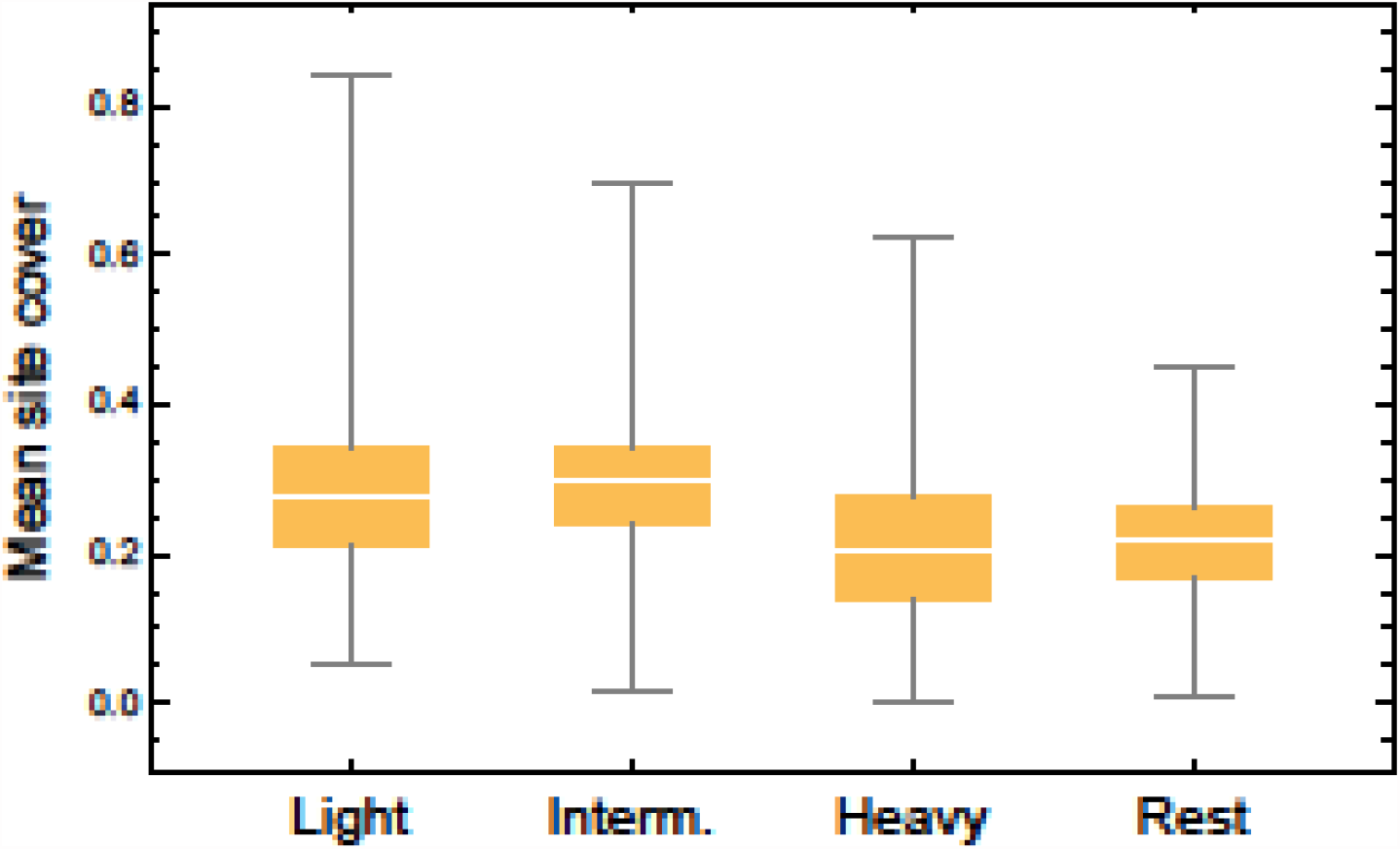
Distribution of the mean site cover of the four different species groups (PFTs).

The classification of each species that was observed in the four grassland types is shown in the Electronic Supplement

### Statistical model

In this study, I consider multi-species (multi-PFTs) pin-point cover data, and it has previously been demonstrated that a reparametrized Dirichlet-multinomial distribution is a suitable candidate distribution to model multi-species pin-point cover data (Damgaard 2015; 2019b; Damgaard et al. 2017). The advantage of using the reparametrized Dirichlet-multinomial distribution is that the observed negative among-species covariation in mean cover are included in the model, and that the degree of within-species spatial aggregation is taken into account and explicitly modelled by a parameter *δ* (Damgaard 2018). If the spatial aggregation, which lead to over-dispersed cover data, had been ignored then the statistical inferences would have been biased.

More specifically, the mean plant cover of a species group *i* is assumed to be a linear model of year (*y*), habitat type (*j*) and site (*k*),

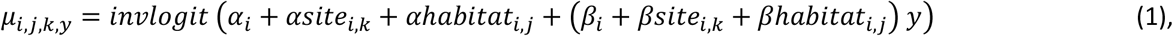

where site and habitat are modelled as random effects. Each random effect, consisting of *n*groups {1, …, *m*, …, *n*}, is modelled as *z*_*m*_ = σ ∈_*m*_, with a standard deviation (σ > 0)and *n*group regressors, ∈_*m*_, that are assumed to have a strong prior probability distribution, ∈_*m*_∼*N*(0,1). For example, *βhabitat*_*i,j*_ = *βhabitat*_σ,*i*_ *βhabitat*_∈,*i,j*_, where the parameter *βhabitat*_σ,*i*_ is positive, and the *n*_*j*_ parameters *βhabitat*_∈,*i,j*_ are assumed to be standard normally distributed. Other location parameters had a weak normally distributed prior probability distribution. Standard deviation parameters were chosen to be uninformative in the positive domain using an exponential prior probability distribution (*λ* = 0.5), and the degree of spatial aggregation had an uninformative uniform prior probability distribution between 0.01 and 0.9.

In the applied general grassland model, there are two hierarchical levels (habitat type and site) with a total of 1469 parameters, and it was not computationally feasible to include plot as a third hierarchical level. This modelling choice was further motivated by the fact that it was unlikely that the location of resampled plots overlapped, and sampling could therefore be assumed to be performed at the site level. Furthermore, an autoregressive component at the plot level did not alter results qualitatively when modeling the change in cover of single heathland habitats, even though the vegetation at these habitats is rather spatially aggregated (Damgaard 2012; Damgaard 2019b).

The model was fitted in MathematicaStan (cmdstan-2.18.0) (Carpenter et al. 2017) using three MCMC chains of 100,000 iterations after a warmup of 100,000 iterations. The STAN model and the Mathematica notebook, including trace plots, are included as electronic supplements.

In order to ensure that the sum of the mean cover parameters did not exceed one when the parameters in the linear model (1) were initialized to zero at the beginning of the MCMC sampling procedure, 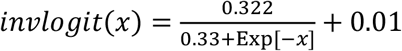 was used instead of transforming the linear model with the standard inverse logit function (Damgaard 2018).

Statistical inferences were based on the marginal posterior probability distributions of parameters and compound parameters, i.e. their 2.5%, 50% and 97.5% percentiles, and the proportion of distribution that is larger than zero, P(>0).

## Results

The fitted model had relatively many parameters, nevertheless, the MCMC iterations demonstrated good mixing properties and the marginal posterior probability distribution of the parameters had regular shapes (see Appendix S4). Therefore, it was concluded that the model fitted the data adequately. The marginal posterior probability distributions of selected parameters and compound parameters are summarized in Tables 1, 2 and 3.

**Table 1.** Summary of the marginal posterior probability distributions of the general parameters in model (1) for the different plant functional types (PFTs).

| Parameter | PFTs | Mean | SD | 2.5% | 50% | 97.5% | P(>0) |
| --- | --- | --- | --- | --- | --- | --- | --- |
| $\alpha$ | Light seeds | 0.050 | 0.335 | -0.669 | 0.060 | 0.747 | 0.601 |
| $\alpha$ | Intermediate | 0.077 | 0.467 | -1.084 | 0.104 | 1.005 | 0.630 |
| $\alpha$ | Heavy seeds | -0.285 | 0.421 | -1.134 | -0.305 | 0.750 | 0.186 |
| $\alpha_{site_{\sigma}}$ | Light seeds | 0.406 | 0.024 | 0.361 | 0.406 | 0.456 | - |
| $\alpha_{site_{\sigma}}$ | Intermediate | 0.474 | 0.027 | 0.423 | 0.473 | 0.529 | - |
| $\alpha_{site_{\sigma}}$ | Heavy seeds | 0.554 | 0.033 | 0.492 | 0.552 | 0.622 | - |
| $\alpha_{habitat_{\sigma}}$ | Light seeds | 0.578 | 0.384 | 0.208 | 0.457 | 1.693 | - |
| $\alpha_{habitat_{\sigma}}$ | Intermediate | 0.873 | 0.732 | 0.268 | 0.644 | 3.636 | - |
| $\alpha_{habitat_{\sigma}}$ | Heavy seeds | 0.759 | 0.625 | 0.242 | 0.581 | 2.684 | - |
| $\beta$ | Light seeds | 0.000 | 0.025 | -0.047 | -0.001 | 0.047 | 0.476 |
| $\beta$ | Intermediate | -0.013 | 0.017 | -0.041 | -0.013 | 0.016 | 0.099 |
| $\beta$ | Heavy seeds | -0.015 | 0.013 | -0.040 | -0.014 | 0.006 | **0.048** |
| $\beta_{site_{\sigma}}$ | Light seeds | 0.024 | 0.006 | 0.011 | 0.024 | 0.034 | - |
| $\beta_{site_{\sigma}}$ | Intermediate | 0.028 | 0.005 | 0.018 | 0.028 | 0.037 | - |
| $\beta_{site_{\sigma}}$ | Heavy seeds | 0.025 | 0.008 | 0.008 | 0.026 | 0.039 | - |
| $\beta_{habitat_{\sigma}}$ | Light seeds | 0.035 | 0.038 | 0.007 | 0.024 | 0.132 | - |
| $\beta_{habitat_{\sigma}}$ | Intermediate | 0.019 | 0.028 | 0.001 | 0.011 | 0.083 | - |
| $\beta_{habitat_{\sigma}}$ | Heavy seeds | 0.014 | 0.022 | 0.000 | 0.008 | 0.065 | - |

**Table 2.**
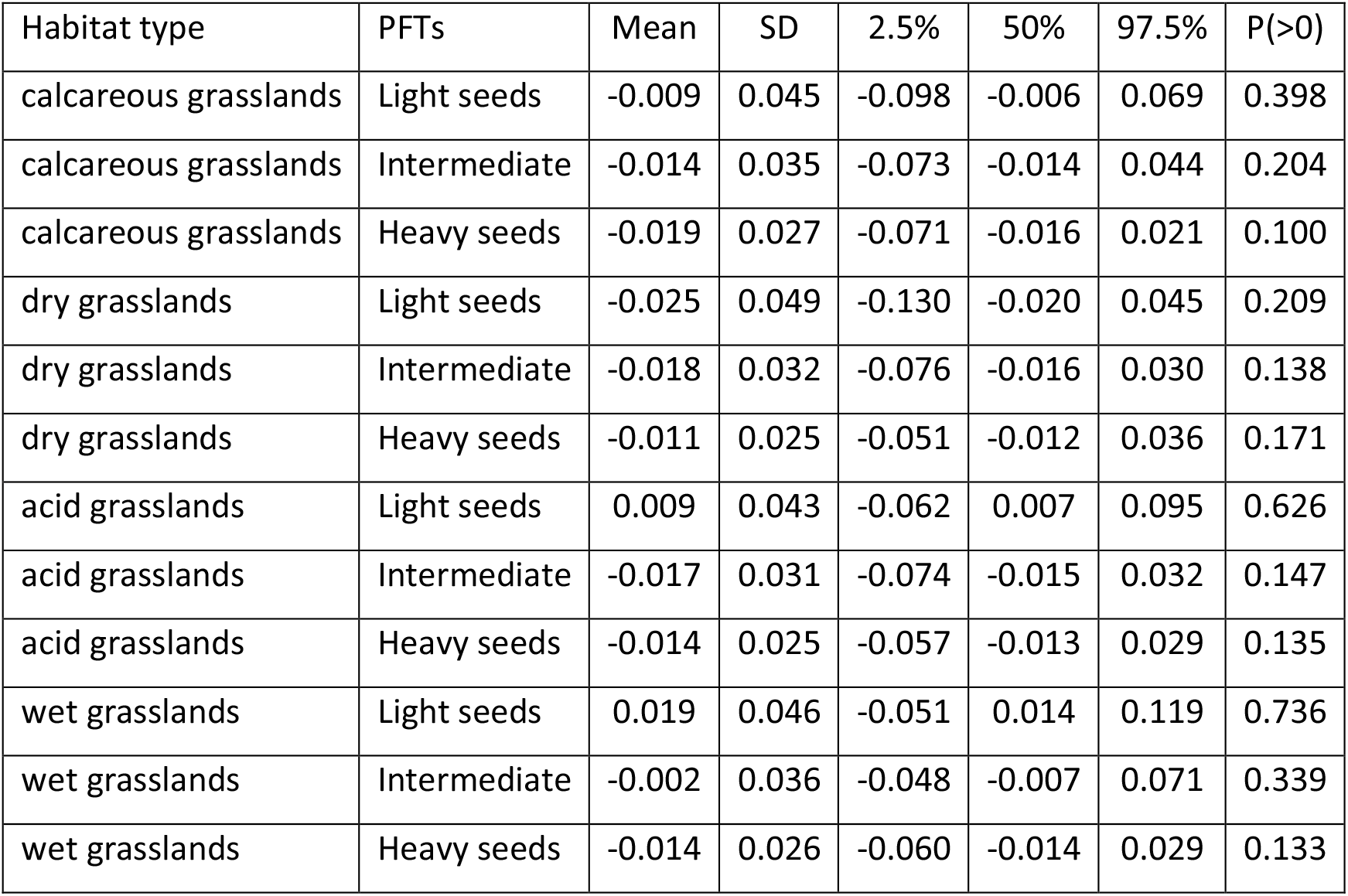
Summary of the marginal posterior probability distribution of the change in cover (*β*_*i*_ + *βhabitat*_σ,*i*_ *βhabitat*_∈,*i,j*_)for the different plant functional types (PFTs) in the different grassland habitat types.

| Habitat type | PFTs | Mean | SD | 2.5% | 50% | 97.5% | P(>0) |
| --- | --- | --- | --- | --- | --- | --- | --- |
| calcareous grasslands | Light seeds | -0.009 | 0.045 | -0.098 | -0.006 | 0.069 | 0.398 |
| calcareous grasslands | Intermediate | -0.014 | 0.035 | -0.073 | -0.014 | 0.044 | 0.204 |
| calcareous grasslands | Heavy seeds | -0.019 | 0.027 | -0.071 | -0.016 | 0.021 | 0.100 |
| dry grasslands | Light seeds | -0.025 | 0.049 | -0.130 | -0.020 | 0.045 | 0.209 |
| dry grasslands | Intermediate | -0.018 | 0.032 | -0.076 | -0.016 | 0.030 | 0.138 |
| dry grasslands | Heavy seeds | -0.011 | 0.025 | -0.051 | -0.012 | 0.036 | 0.171 |
| acid grasslands | Light seeds | 0.009 | 0.043 | -0.062 | 0.007 | 0.095 | 0.626 |
| acid grasslands | Intermediate | -0.017 | 0.031 | -0.074 | -0.015 | 0.032 | 0.147 |
| acid grasslands | Heavy seeds | -0.014 | 0.025 | -0.057 | -0.013 | 0.029 | 0.135 |
| wet grasslands | Light seeds | 0.019 | 0.046 | -0.051 | 0.014 | 0.119 | 0.736 |
| wet grasslands | Intermediate | -0.002 | 0.036 | -0.048 | -0.007 | 0.071 | 0.339 |
| wet grasslands | Heavy seeds | -0.014 | 0.026 | -0.060 | -0.014 | 0.029 | 0.133 |

**Table 3.** Summary of the marginal posterior probability distribution of the degree of spatial aggregation in the different grassland habitat types.

| Habitat type | Mean | SD | 2.5% | 50% | 97.5% |
| --- | --- | --- | --- | --- | --- |
| calcareous grasslands | 0.128 | 0.007 | 0.116 | 0.128 | 0.142 |
| dry grasslands | 0.074 | 0.002 | 0.071 | 0.074 | 0.077 |
| acid grasslands | 0.077 | 0.002 | 0.073 | 0.077 | 0.080 |
| wet grasslands | 0.132 | 0.003 | 0.126 | 0.132 | 0.139 |

Across the four grassland habitat types, there was community selection towards lighter seeds. The cover of heavy seeds decreased significantly during the eight-year period (Table 1: heavy seeds, *P*(*β* > 0)= 0.048) and the cover of intermediate seeds showed a tendency towards a decrease (Table 1: intermediate seeds, *P*(*β* > 0)= 0.099). The probability of both these observed decreases in cover should occur by chance is 0.005. However, the selection towards lighter seeds was not significant when considering each of the four grassland habitats separately (Table 2), possibly due to reduced statistical power. The distribution of the mean site cover at the four grassland habitat types over the years is shown in Fig. 3.

**Fig. 3.**
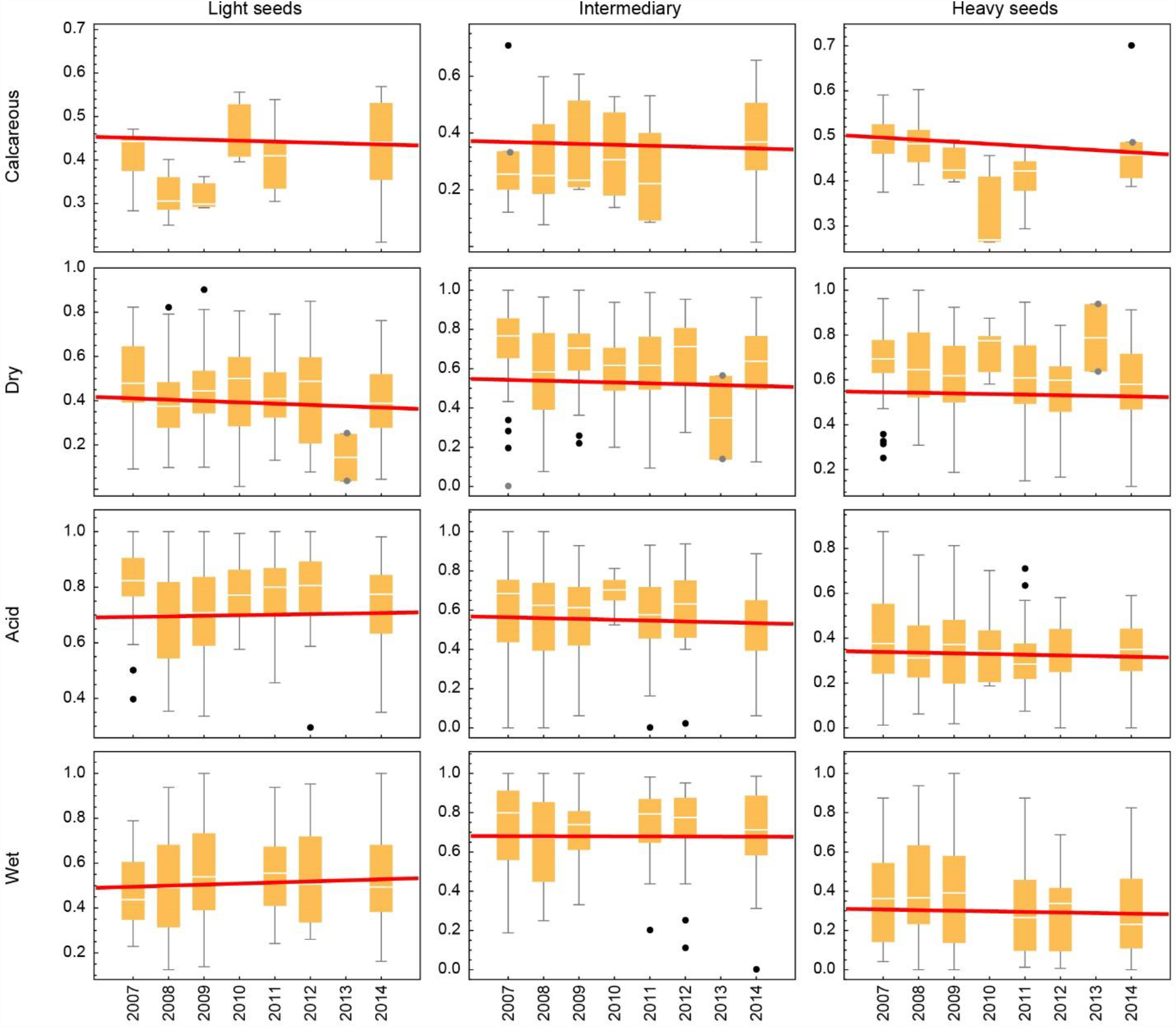
Distribution of the mean site cover at the four grassland habitat types over the years. The estimated slopes are illustrated approximately with the red lines. Note that the considerable among year variation mainly is due to an irregular sampling design.

In order to examine possible patterns of the among-site variation the mean of the random effects for the different aggregated species groups, *E*(*βsite*_*i,k*_)were compared to information on whether signs of grazing were observed in each plot during the years, which was assumed to be a proxy for management intensity. However, no significant patterns were observed (Appendix S4).

The estimated standard deviations of the yearly change among sites were approximately of the same size as the standard deviations among habitats (Table 1), which indicates that there is a sizeable variation in the yearly change among sites of the same grassland habitat type.

The standard deviations of the posterior probability distributions of the slope parameters that measure the yearly change in the cover of the PFTs were approximately 0.02 on the logit scale (Table 1). This width of the posterior probability distribution indicates that the statistical power of the analysis is moderate and, in connection with the mostly non-significant changes in cover, it is concluded that trait selection for seed mass in Danish grasslands over the eight-year period generally is week.

The spatial aggregation of the FPTs was significantly higher in calcareous grasslands and wet grasslands than in the two other grassland habitat types (Table 3).

## Discussion

Across the four grassland habitat types, there was selection at the community level towards lighter seeds during an eight-year period. I am not aware of other studies that have investigated selection on seed mass at the community level using time-series data from a large number of grassland sites. Unfortunately, a general theory on community level selection on seed mass in a changing environment and motivated hypotheses on what to expect under different circumstances are underdeveloped (although see e.g. Larios and Venable 2018). To further generate such hypotheses, possible patterns between the among-site variation and observed signs of grazing were examined, but did not reveal any significant patterns. It may be speculated that the observed community selection towards lighter seeds may be due to reduced abiotic stress in more productive grasslands, since it is known that heavy seeds are more likely to tolerate stressful conditions early in plant development (Coomes and Grubb 2003; Larios et al. 2014; Westoby et al. 2002).

There has been an extensive exploration of the role of different plant traits, including the role of seed mass, to characterize different plant life history strategies and how the different strategies are more or less suited to different environments (e.g. Garnier et al. 2016; Westoby et al. 2002). For example, annual plants generally produce lighter seeds than biennials or perennial plants, tall-statured plants produce bigger seeds than small plants (e.g. Thomson et al. 2011), and large seeds are considered an adaptation to stable environments (e.g. Salisbury 1975). It is indeed possible that the changing environment has caused the observed decline in large-seeded species in Danish grasslands. However, more time-series studies are needed in order to understand the relevant ecological processes rather than relying on the commonly performed space-for-time approach (Damgaard 2019a).

The ultimate goal of a community ecological investigation is to understand the underlying causes of an observed pattern, which would enable us to make credible ecological predictions. However, a specific ecological pattern may be caused by several different processes and involve several important contingencies (Vellend 2010), and the relationship between process and pattern may be influenced by time lags of unknown duration (Svenning and Sandel 2013). Consequently, it will not always be possible to link an observed ecological pattern to its underlying causes, and this is especially true if the investigation is performed at a relatively large spatial scale, where it is a more demanding task to monitor all the possible causal factors (Vellend 2010).

The studied vegetation plots were not real permanent plots, but were only resampled with GPS-uncertainty. However, if there were access to permanent plot time-series cover data, it would have been interesting to explore more sophisticated selection models, e.g. whether the observed trait selection could be partitioned into direct selection and selection that is mediated by interspecific interactions (Damgaard 2016; Pedersen et al. 2019).

In order to model the sampling variance correctly and avoid possible model bias (Clark 2016), the continuous variable, seed mass, is used for grouping plant species into PFTs, and the aggregated cover of the PTFs is then treated as the dependent variable. This method is relatively conservative compared to treating the CWMs of seed mass as the dependent variable. However, in order to ensure that the reported results are representative and may be generalized to other areas, it is of utmost importance that the sampling variance is modelled correctly.

An alternative method of analyzing plant trait selection is to correlate trait values with estimated demographic parameters or change in abundance for individual species (Garnier et al. 2018; Herben et al. 2019; Timmermann et al. 2015). However, it may be a problem that typically only point estimates of the plant ecological success are used without considering the covariance between plant abundance and trait variation (Clark 2016). Furthermore, since many plant traits are correlated (e.g. Reich 2014), it is problematic to infer from observed correlations to causal mechanisms. Especially since the studied plant traits often are the traits that are most readily available in trait databases.

In this study, plant abundance is measured by plant cover using the pin-point method, which readily allows the aggregation of single cover into the cover of PFTs at the pin level. As demonstrated elsewhere, it is important to take the spatial aggregation of plant species into account when modelling plant cover (Damgaard 2012; 2013; Damgaard and Irvine 2019), and these results have been generalized into a multi-species setting, where it is recommended to analyze multispecies or PFTs pin-point cover data in a reparametrized Dirichlet-multinomial distribution (Damgaard 2015; 2018). A further advantage of modelling trait selection by the change in the cover of PFTs rather than the change in the continuous CWMs is that the class of missing values, i.e. the plant species with no or incomplete information on the trait values, is clearly defined (in this study as the rest group). Additionally, more complicated selection models than directional selection may be readily tested, e.g. stabilizing selection (the intermediate PFT is positively selected) and disruptive selection (the intermediate PFT is negatively selected).

## Data Accessibility statement

The used cover data may be found in Appendix S1

## Author Contribution

CD analyzed the data and wrote the manuscript

## Electronic Supplements

All Appendices may be downloaded from https://osf.io/n7mu3/

Appendix S1: Cover data

Appendix S2: Species list (Excel file)

Appendix S3: STAN model (text file)

Appendix S4: Mathematica notebook with trace plots and simulated posterior probability distributions etc.

## References

Carpenter B, Gelman A, Hoffman MD, Lee D, Goodrich B, Betancourt M, Brubaker M, Guo J, Li P, Riddell A. 2017. Stan: A probabilistic programming language. 2017. 76(1):32.

Clark JS. 2016. Why species tell more about traits than traits about species: Predictive analysis. Ecology. 97(8):1979–1993.

Coomes DA, Grubb PJ. 2003. Colonization, tolerance, competition and seed-size variation within functional groups. Trends Ecol Evol. 18(6):283–291.

Damgaard C. 2009. On the distribution of plant abundance data. Ecol Inform. 4:76–82.

Damgaard C. 2012. Trend analyses of hierarchical pin-point cover data. Ecology. 93:1269–1274.

Damgaard C. 2013. Hierarchical and spatially aggregated plant cover data. Ecol Inform. 18:35–39.

Damgaard C. 2015. Modelling pin-point cover data of complementary vegetation classes. Ecol Inform. 30:179–184.

Damgaard C. 2016. Empirical modelling of trait selection by partitioning selection into direct selection and selection that is mediated by interspecific interactions. bioRxiv.

Damgaard C. 2018. The joint distribution of pin-point plant cover data: A reparametrized dirichlet - multinomial distribution. arXiv e-prints. [accessed August 01, 2018]https://ui.adsabs.harvard.edu/\#abs/2018arXiv180804582D.

Damgaard C. 2019a. A critique of the space-for-time substitution practice in community ecology. Trends Ecol Evol. 34(5):416–421.

Damgaard C. 2019b. Spatio-temporal structural equation modeling in a hierarchical bayesian framework: What controls wet heathland vegetation? Ecosystems. 22:152–164.

Damgaard C. 2021. Indication of a reduction in the cover of thin-leaved plants in danish grasslands over an eight-year period. Journal of Vegetation Science. n/a(n/a):e12982.

Damgaard C, Hansen RR, Hui FKC. 2020. Model-based ordination of pin-point cover data: Effect of management on dry heathland. Ecol Inform. 60:101155.

Damgaard C, Irvine KM. 2019. Using the beta distribution to analyze plant cover data. J Ecol. 107:2747–2759.

Damgaard C, Nielsen KE, Strandberg M. 2017. The effect of nitrogen deposition on the vegetation of wet heathlands. Plant Ecol. 218(4):373–383.

Ellermann T, Bossi R, Nygaard J, Christensen J, Løfstrøm P, Monies C, Grundahl L, Geels C, Nielsen IE, Poulsen MB. 2019. Atmosfærisk deposition 2017. Aarhus University.

EU. 2013. Interpretation manual of european union habitats.. Bruxelles: European Commission, DG Environment, Nature and Biodiversity.

Garnier E, Fayolle A, Navas M-L, Damgaard C, Cruz P, Hubert D, Richarte J, Autran P, Leurent C, Violle C. 2018. Plant demographic and functional responses to management intensification: A long-term study in a mediterranean rangeland. J Ecol. 106(4):1363–1376.

Garnier E, Navas ML, Grigulis K. 2016. Plant functional diversity. Organism traits, community structure, and ecosystem properties. Oxford, UK: Oxford University Press.

Herben T, Hadincová V, Krahulec F, Pecháčková S, Skálová H. 2019. Two dimensions of demographic differentiation of species in a mountain grassland community: An experimental test. Functional Ecology. 33(8):1514–1523.

Hui FKC, Taskinen S, Pledger S, Foster SD, Warton DI. 2015. Model-based approaches to unconstrained ordination. Methods in Ecology and Evolution. 6(4):399–411.

Jansen PA, Bartholomeus M, Bongers F, Elzinga JA, Ouden J, van den Wieren SE. 2002. The role of seed size in dispersal by a scatter-hoarding rodent. In: Levey DJ, Silva WR, Galetti M, editors. Seed dispersal and frugivory: Ecology, evolution and conservation third international symposium-workshop on frugivores and seed dispersal, são pedro, brazil, 6–11 august 2000. CABI.

Kleyer M, Bekker RM, Knevel IC, Bakker JP, Thompson K, Sonnenschein M, Poschlod P, Van Groenendael JM, Klimeš L, Klimešová J et al. 2008. The leda traitbase: A database of life-history traits of the northwest european flora. J Ecol. 96(6):1266–1274.

Larios E, Búrquez A, Becerra JX, Lawrence Venable D. 2014. Natural selection on seed size through the life cycle of a desert annual plant. Ecology. 95(11):3213–3220.

Larios E, Venable DL. 2018. Selection for seed size: The unexpected effects of water availability and density. Functional Ecology. 32(9):2216–2224.

Levy EB, Madden EA. 1933. The point method of pasture analyses. New Zealand Journal of Agriculture. 46:267–279.

Lindquist B. 1931. Den skandinaviska bokskogens biologi. Svenska Skogsvårdsföeningens Tidskrift. 3:179–485.

Maron JL, Hahn PG, Hajek KL, Pearson DE. 2021. Trade-offs between seed size and biotic interactions contribute to coexistence of co-occurring species that vary in fecundity. J Ecol. 109(2):626–638.

Muller-Landau HC. 2010. The tolerance–fecundity trade-off and the maintenance of diversity in seed size. Proc Natl Acad Sci. 107(9):4242.

Nielsen KE, Bak JL, Bruus M, Damgaard C, Ejrnæs R, Fredshavn JR, Nygaard B, Skov F, Strandberg B, Strandberg M. 2012. Naturdata.Dk - danish monitoring program of vegetation and chemical plant and soil data from non-forested terrestrial habitat types. Biodiversity & Ecology 4:375.

Nygaard B, Damgaard C, Nielsen KE, Bladt J, Ejrnæs R. 2016. Habitatdirektivets naturtyper. Aarhus Universitet, DCE – Nationalt Center for Miljø og Energi.

Nygaard B, Ejrnæs R, Baattrup-Pedersen A, Fredshavn J. 2009. Danske plantesamfund i moser og enge – vegetation, økologi, sårbarhed og beskyttelse. Aarhus: DMU.

Pedersen RØ, Bonis A, Damgaard C. 2019. A nonlinear bayesian model of trait selection forces. Ecological Modelling. 393:107–119.

Poschlod P, Bakker JP, Kahmen S. 2005. Changing land use and its impact on biodiversity. Basic and Applied Ecology. 6(2):93–98.

Reich PB. 2014. The world-wide ‘fast–slow’ plant economics spectrum: A traits manifesto. J Ecol. 102(2):275–301.

Salisbury EJ. 1975. The survival value of modes of dispersal. Proceedings of the Royal Society of London Series B Biological Sciences. 188(1091):183–188.

Svenning J-C, Sandel B. 2013. Disequilibrium vegetation dynamics under future climate change. Am J Bot. 100:1–21.

Thomson FJ, Moles AT, Auld TD, Kingsford RT. 2011. Seed dispersal distance is more strongly correlated with plant height than with seed mass. J Ecol. 99(6):1299–1307.

Timmermann A, Damgaard C, Strandberg MT, Svenning J-C. 2015. Pervasive early 21st-century vegetation changes across danish semi-natural ecosystems: More losers than winners and a shift towards competitive, tall-growing species. J Appl Ecol. 52(1):21–30.

Vellend M. 2010. Conceptual synthesis in community ecology. The Quarterly Review of Biology. 85(2):183–206.

Westoby M, Falster DS, Moles AT, Vesk PA, Wright IJ. 2002. Plant ecological strategies: Some leading dimensions of variation between species. Annual Review of Ecology and Systematics. 33(1):125–159.

